# Orthogonal Force Balance Between Contractility and Shear Stress Governs Podocyte Dynamics

**DOI:** 10.64898/2026.01.22.701159

**Authors:** Shumeng Jiang, Pongpratch Puapatanakul, Chengqing Qu, Yin-Yuan Huang, Yuxuan Huang, Kaye E. Brathwaite, Carmen M. Halabi, Jeffrey H Miner, Guy M. Genin, Hani Y Suleiman

## Abstract

Maintenance of tissue barriers under mechanical stress represents a fundamental biological challenge across organ systems. In the kidney, podocyte cells withstand highly variable hemodynamic forces while preserving a tensioned, nanostructured filtration barrier. Dysregulation of this barrier leads to significant pathology, but the mechanical principles underlying homeostasis of cells against flow of filtrates have not yet been identified. Here, we uncover a counterintuitive mechanical homeostasis mechanism whereby podocyte attachment depends on a dynamic balance between external fluid shear stress and internal cellular contractility. Integrated biomechanical modeling and experiment reveal a previously unrecognized mechanosensing circuit that optimizes integrin distribution at foot process peripheries. Our mathematical framework for cell-matrix adhesion stability reveals, surprisingly, that reducing blood pressure can worsen outcomes when cell contractility is impaired, contrary to clinical belief that lowering blood pressure universally benefits cellular adhesion and kidney function. We validated this principle through mouse models with manipulated blood pressure and myosin inhibition, demonstrating that concurrent reduction of both shear stress and contractility worsens podocyte injury and proteinuria. Super-resolution microscopy confirms our predicted integrin redistribution patterns under these mechanical perturbations. These findings establish a fundamental mechanobiological principle applicable beyond nephrology, and suggest potential treatment pathways targeting non-equilibrium steady states.

## Introduction

The kidney glomerulus balances a fundamental mechanobiological paradox: its podocytes must maintain nanometer-scale adhesions while filtering 180 liters of plasma daily under continuous hemodynamic stress ^1^. The podocyte, with its elaborate foot process architecture, forms the final layer of the glomerular filtration barrier through interdigitating structures that create slit diaphragms only 30-40 nm wide ^2–4^. These terminally differentiated cells cannot regenerate ^5^ making their loss into urine (podocyturia) a pathway towards irreversible chronic kidney disease ^1,5–10^. Both loss of proteins to the urine (proteinuria) and podocyturia are associated with hypertension and progression to end-stage kidney disease with increased mortality risk ^11,12^.

Clinically, this mechanical complexity manifests as a therapeutic paradox that challenges our fundamental understanding of podocyte adhesion. While reducing blood pressure effectively decreases proteinuria, podocyte loss continues even with optimal blood pressure control, and lowering systemic blood pressure does not normalize kidney structure ^13–21^. Proteinuria often increases in patients experiencing low blood pressure from systemic shock, with disruption of the glomerular filtration barrier evident ^22,23^, and with mean arterial pressures below a limit exacerbating glomerular injury ^24^. A clue to resolving this mystery is that mechanical force regulation by cells is clearly involved: angiotensin II blockade proves more effective than blood pressure reduction alone in stabilizing glomeruli and reducing podocyte loss ^16–18^, suggesting that cellular contractility and fluid forces interact in ways we do not yet understand. We therefore explored the mechanobiological mechanisms underlying this disconnect that lowering mechanical stress on the glomerulus does not prevent cellular detachment.

While the biochemical factors governing podocyte adhesion and the cytoskeletal proteins essential for foot process formation are well characterized, their mechanical regulation remains unclear ^25,26^. Podocytes experience dynamic mechanical challenges including changes in capillary width, pressure differences, and basement membrane stiffness throughout life and injury ^27–31^, factors known to influence podocyte behavior *in vitro* ^32–34^. The podocyte cytoskeleton is essential for responding to these challenges ^35,36^. Angiotensin II blockade (ACE inhibitors or ARBs) are more effective than blood pressure reduction alone in stabilizing glomeruli and reducing ongoing podocyte loss ^16–18^, indicating that there is an interaction between the effects of cytoskeletal contraction and intraglomerular fluid pressure and flow in the regulation of podocyte attachment.

Although extensive research has elucidated how environmental stiffness and internal cellular forces affect adhesions across diverse cell types, the unique challenge of maintaining adhesion under perpendicular hydrodynamic filtering forces has never been addressed ^37–40^. We hypothesized that podocyte stability requires not simply minimal mechanical stress, but rather a balance between orthogonal forces: external vertical shear stress from filtration must balance against internal horizontal contractile forces required for adhesion. By developing a mathematical model integrating these perpendicular mechanical forces with podocyte cytoskeletal architecture, and testing predictions in a mouse model, we reveal how this delicate force balance governs filtration barrier integrity. Our findings explain why reducing blood pressure alone proves insufficient to preserve podocyte attachment and identify optimal mechanical regimes where both proteinuria and podocyturia can be attenuated, suggesting a new framework for therapeutic intervention in chronic kidney disease.

## Results

### Concurrent reduction of blood pressure and podocyte contractility causes catastrophic barrier failure

To begin assessing how mechanical forces affect podocyte attachment, we challenged the conventional therapeutic approach of reducing blood pressure in kidney disease. We developed a mouse model with bidirectionally tunable blood pressure and glomerular filtration rate (GFR), combined with pharmacological inhibition of podocyte contractility (Fig. 1a). Through AAV-mediated renin expression, we achieved a 20% increase in systolic and diastolic blood pressure, while combination antihypertensive therapy (losartan and prazosin) ^41–43^ produced a 20% reduction (Fig. 1b). These manipulations translated to corresponding changes in GFR: increased blood pressure elevated GFR by 30% specifically at day 5 post-treatment (Long-term hypertension injures kidney and eventually reduce GFR (Supplementary Fig. 1)), while antihypertensive therapy reduced GFR by 25% (Fig. 1c), confirming that our interventions successfully modulated shear stress at the filtration barrier.^44^

**Figure 1.**
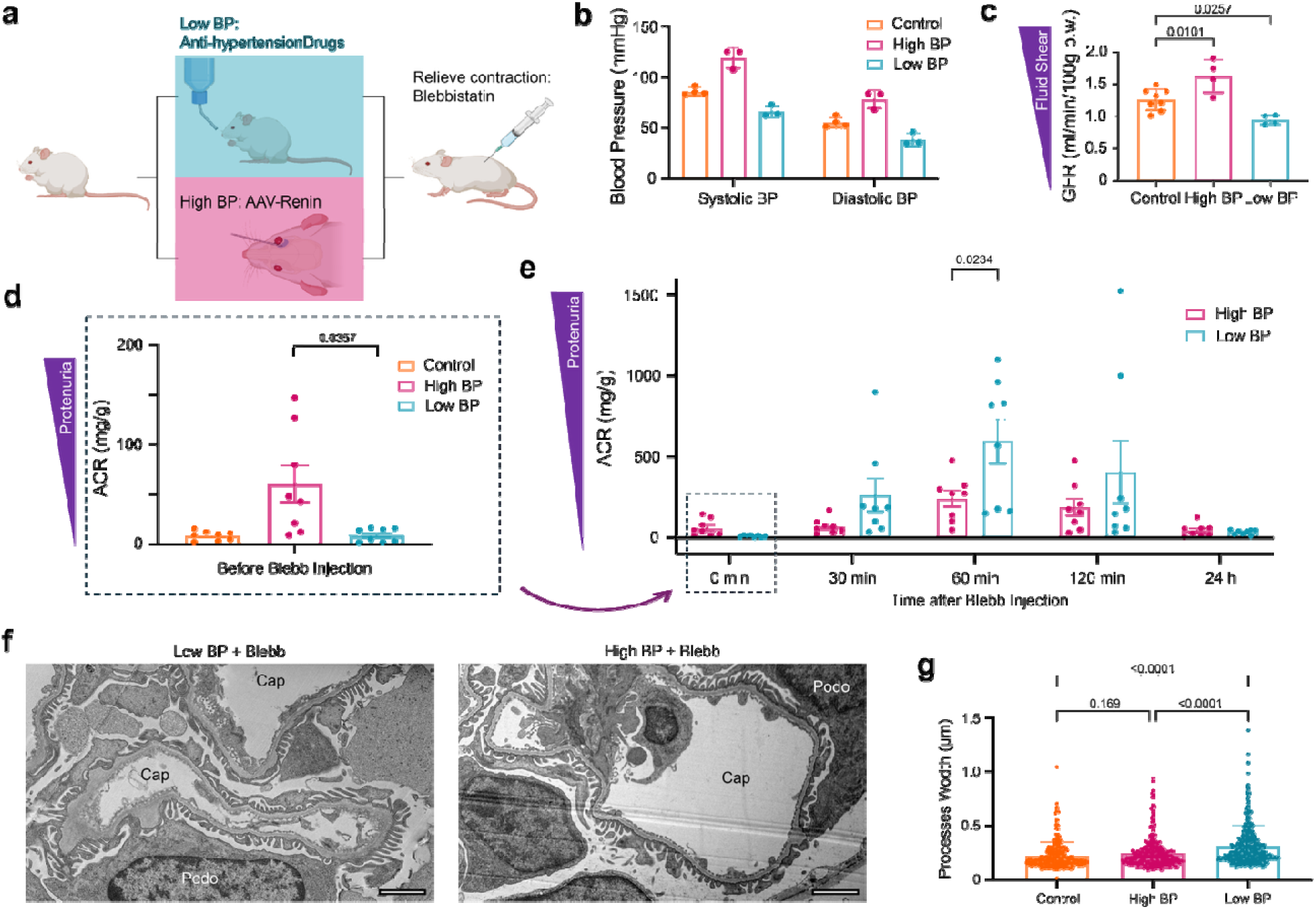
Concurrent reduction of blood pressure and podocyte contractility causes catastrophic glomerular barrier failure. **a,** Schematic of experimental design showing bidirectional blood pressure manipulation through AAV-renin injection or antihypertensive drugs combined with blebbistatin injection to inhibit podocyte contractility. **b,** Blood pressure (BP) manipulation achieved through experimental interventions. AAV-renin injection increases systolic and diastolic BP by 20% after one month, while antihypertensive therapy (losartan + prazosin) reduces BP by 20%. Data shown as mean ± SD, n = 3 mice per group. **c,** Glomerular filtration rate (GFR) changes correspond to BP alterations. High BP increases GFR by 30% at day 5 post-AAV injection, while antihypertensive treatment reduces GFR by 25%, confirming successful modulation of shear stress at the filtration barrier. **e,** Paradoxical injury response: Low blood pressure (BP) mice develop severe proteinuria (ACR > 500 μg/mg) 60 minutes after contractility inhibition with blebbistatin, significantly exceeding the response in high BP mice (*p* < 0.05), **d**, despite showing no baseline proteinuria. Data shown as mean ± SD, *n* = 8 mice per group. **f-g,** Transmission electron microscopy validates predicted mechanical failure. Low BP mice with inhibited contractility show foot process effacement (mean width 420 ± 120 nm vs 210 ± 45 nm control, *p* < 0.001) and characteristic GBM thickening and undulation (arrows) 60 min post-blebbistatin. High BP mice maintain foot process architecture despite contractility inhibition. Scale bar: 2 μm.

Paradoxically, when we inhibited podocyte contractility with the myosin II inhibitor blebbistatin ^45^, known to disrupt podocyte stress fibers and contractile “sarcomere-like structures” ^46^, mice with reduced blood pressure developed the most severe kidney injury. Before blebbistatin injection, low blood pressure mice showed no proteinuria (albumin-to-creatinine ratio (ACR) < 30 μg/mg), while high blood pressure mice exhibited mild albuminuria (ACR ∼100 μg/mg), consistent with hypertensive injury (Fig. 1d). However, 60 minutes after blebbistatin administration (10 mg/kg intraperitoneally) ^47,48^, low blood pressure mice developed severe proteinuria (ACR > 500 μg/mg), significantly exceeding the response in high blood pressure mice (ACR ∼200 μg/mg, *p* < 0.01) (Fig. 1e). This transient but dramatic barrier failure, returning to baseline by 24 hours, revealed that reducing both external shear stress and internal contractility simultaneously caused catastrophic podocyte dysfunction.

Ultrastructural analysis confirmed the severity of injury in low blood pressure mice with inhibited contractility. Transmission electron microscopy (TEM) performed 60 min post-blebbistatin injection revealed extensive foot process effacement, with mean foot process width increasing from 210 ± 45 nm to 420 ± 120 nm (*p* < 0.001), accompanied by characteristic glomerular basement membrane thickening and undulation (Fig. 1f, g). High blood pressure mice showed minimal structural changes despite blebbistatin treatment, suggesting that maintained shear stress partially compensates for lost contractility. These findings challenge the prevailing clinical paradigm that reducing mechanical stress universally protects podocytes, instead revealing that podocyte attachment requires a critical balance between opposing mechanical forces. To understand this counterintuitive mechanism, we developed a mathematical framework integrating the unique mechanical environment of the filtration barrier.

### Mathematical modeling reveals orthogonal force balance governs podocyte attachment

To understand this paradoxical response, we developed a biomechanical model incorporating the unique three-dimensional force environment of podocyte foot processes. Unlike traditional cell-matrix adhesion models where forces act parallel to the substrate ^49,50^, podocytes experience orthogonal mechanical challenges: vertical shear stress from glomerular filtration opposes horizontal contractile forces generated within the cell body and transmitted through actin cables (Fig. 2a).

**Figure 2.**
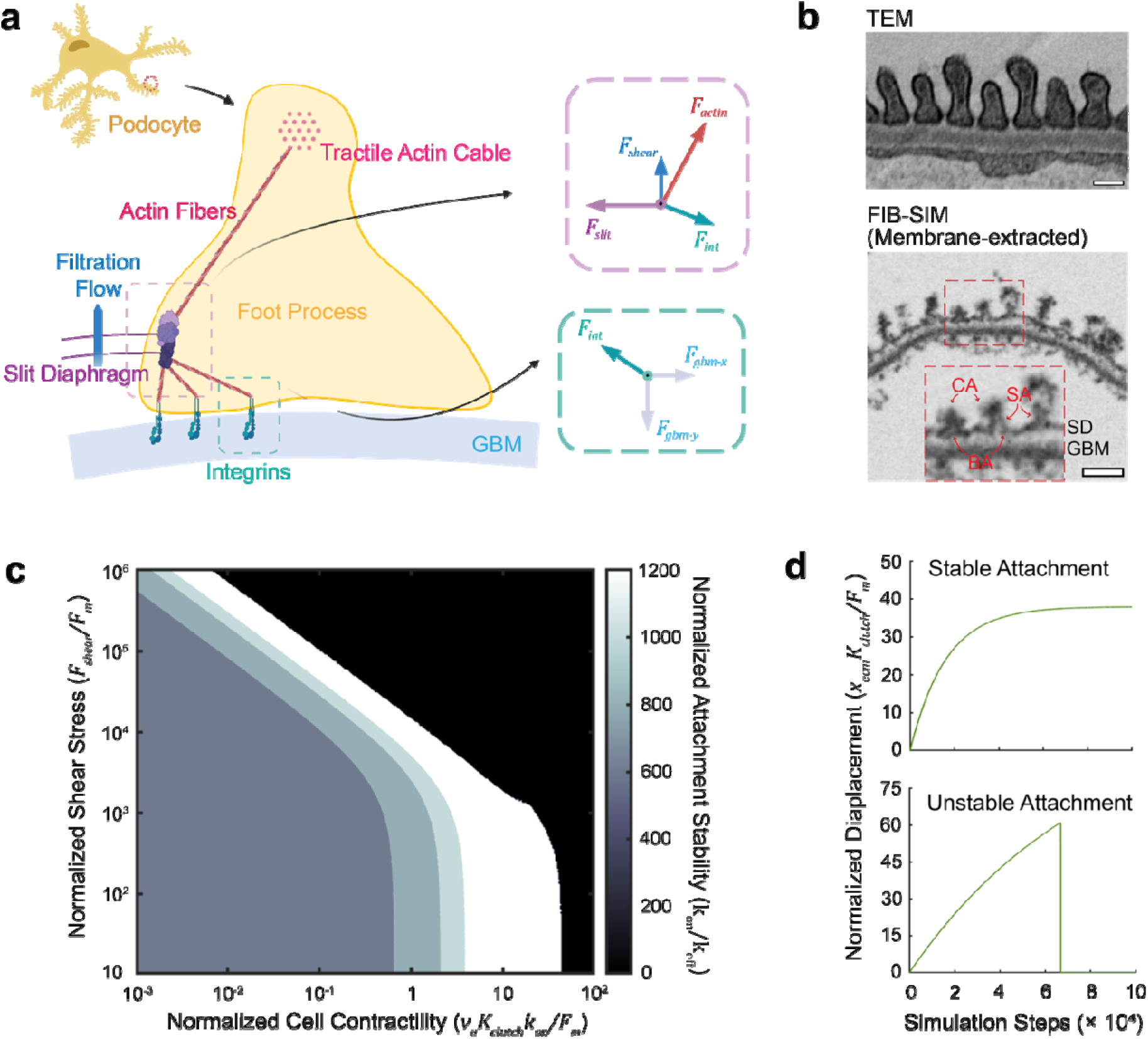
Mathematical modeling predicts loss of orthogonal force balance led to the unstable foot processes attachment and kidney injury. **a**, Mathematical model of orthogonal force balance in podocyte foot processes. Vertical shear stress from glomerular filtration () opposes horizontal contractile forces (F_actin) transmitted through central actin cables. Forces converge at slit diaphragm junctions and are balanced by integrin-mediated adhesion to the glomerular basement membrane (GBM). **b,** TEM and FIB-SIM imaging of healthy or membrane extracted mouse kidney tissue, both show existence of central actin cable (CA), side actin (SA) and bottom actin (BA) in foot processes. Scale bar: 200 nm. **c,** Heatmap showing predicted foot process attachment stability across varying shear stress and cell contractility parameters. Color indicates integrin bond stability (ratio). White region indicates stable adhesion; blue indicates unstable attachment leading to detachment. Red lines demarcate physiological boundaries. **d,** Representative displacement curves during Monte Carlo simulations demonstrating three mechanical regimes: stable adhesion with balanced forces (green), unstable progression to detachment with reduced forces (blue), and catastrophic detachment with combined force reduction (red).

An obvious consequence of the free body diagrams is that a structure must exist that connects the actin/nephrin junction to the actin-associated integrin proteins bound to the basement membrane. Although such structures have not been reported in the literature, we repeated earlier membrane extraction and electron microscopy protocol ^51^ to search for these. This imaging indeed revealed dense cytoskeleton structures exist on the center top of the foot processes, probably contractile central actin cables (CA) extending from the cell body, as well as at the side (SA) and the bottom (BA) of the foot processes, probably actin linkage between central actin cable, slit diaphragm and the GBM (Fig. 2b). affirming the structural basis for our mechanical framework (Fig. 2a).

We adapted a molecular clutch model ^49,50^ to account for these orthogonal forces and provide testable predictions of our structural model. In our formulation, contractile forces from the podocyte cell body generate horizontal tension (F_contract_) transmitted through central actin cables, while filtration creates vertical shear stress (r_shear_) acting perpendicular to the adhesion plane. These forces converge at critical junction points where actin cables connect to both the slit diaphragm (maintaining lateral foot process connections) and integrin clusters (securing GBM attachment). In the model, the net force balance at these junctions determined adhesion stability. Our model revealed that these orthogonal forces create a mechanical circuit where each component stabilizes the other. Contractile forces pre-tension the cytoskeleton, enabling it to resist perpendicular shear deformation, while shear stress maintains foot process architecture that optimizes contractile force transmission.

We parameterized integrin dynamics using established force-dependent binding kinetics for integrin αVβ3 (Supplementary Fig 2) ^52^, which shares a degree of homology with the podocyte-specific integrin α3β1, including phosphorylation of focal adhesion kinase.^53–56^ Monte Carlo simulations revealed a non-monotonic relationship: both excessive and insufficient mechanical loading destabilized adhesion (Fig. 2c). The model predicted a mechanical region of homeostasis where combined forces optimize integrin stability: too little total force fails to activate mechanosensitive adhesion strengthening, while excessive force overwhelms molecular bonds. The model predicted that simultaneous reduction of both force components would cause more severe failure than reducing either alone, as the system loses both mechanical activation and structural pre-tension (Fig. 2c). This prediction matches our experimental observation that concurrent blood pressure reduction and contractility inhibition causes catastrophic proteinuria, suggesting a mechanistic explanation for why conventional antihypertensive therapy alone cannot prevent podocyte loss.

### Force balance requirements predict a mechanical steady-state regime that enables stable attachment

Having established the orthogonal force framework, we next explored how varying mechanical parameters across the clinically achievable range affects podocyte attachment stability. We mapped the parameter space by varying shear stress from blood filtration, contractility of actin fibers from myosin functionality in the podocyte cell body, and the stiffness of the GBM. By generating a heatmap to color code the overall stability, we predicted how the system would respond to changes in these parameters.

Shear stress and cell contractility both showed similar effects on integrin stability. An optimum range of each parameter was linked to the highest integrin stability, while excessive or insufficient stress resulted in unstable attachment (Fig. 2c). Lowered stress was associated with decreased integrin stability, while increased stress led to complete detachment of the foot processes, indicated by all integrin bonds breaking simultaneously during the simulation before reaching homeostasis in the system (Fig. 2d).

Synergistic effects between the two parameters were evident, with higher cell contractility reducing the required shear stress to maintain system stability. This compensatory relationship creates a mechanical circuit where each force component can partially substitute for the other. However, this redundancy also reveals a vulnerability: when both shear stress and contractility are reduced simultaneously, the system loses this compensatory mechanism.

The interplay between cytoskeletal contractility, blood pressure, and GBM stiffness, revealed that the stability of the system was predominantly influenced by stresses like shear stress or cell contractility. However, increased GBM stiffness appeared to contribute positively to maintaining stable attachment at low stress, as indicated by an expanded tolerance to varied stress situations (Supplementary Fig. 3).

Cell contractility in the podocyte appears to act as a compensatory mechanism, adjusting to changes in the mechanical microenvironment by accommodating alterations in shear stress and maintaining overall system stability. Disrupting the contractility in the podocyte may render the foot processes more susceptible to changes in shear stresses, especially when shear stress decreases, and there is no cell contractility to compensate for it. This prediction directly aligned with our experimental observation that the combination of decreased shear stress and inhibited contractility in podocytes would lead to more severe injury and proteinuria than either condition alone.

### Mechanical disequilibrium drives podocyte detachment and barrier disruption

To validate the predicted mechanical failure under combined force reduction, we examined kidney tissue samples collected 60 minutes after blebbistatin injection to visualize podocyte and glomerular injury. Ultrathin sectioning and TEM were applied to observe foot process morphology and identify potential foot process effacement in the glomerulus.

The low blood pressure mice that exhibited higher proteinuria showed clear signs of injury including thickened and wavy glomerular basement membrane compared to mice with no blood pressure alteration or high blood pressure (Fig. 1f). Quantitative analysis revealed that mean foot process width increased significantly in the low blood pressure group, suggesting early-stage injury-induced foot process effacement (Fig. 1g). These ultrastructural changes directly corresponded to the severity of proteinuria observed in the functional studies.

In contrast, high blood pressure mice showed minimal structural changes despite blebbistatin treatment, suggesting that maintained shear stress partially compensates for lost contractility. This differential response between low and high blood pressure conditions under contractility inhibition provides structural evidence for the mechanical compensation predicted by our model.

The observation of GBM thickening within certain area after just one hour of blebbistatin injection in low blood pressure mice indicates that podocyte contractility might be essential for proper GBM organization. This rapid structural disruption suggests that the balance between external shear forces and internal contractility not only maintains podocyte adhesion but also regulates basement membrane architecture. The waviness observed in the GBM further suggests loss of the normal tension maintained by properly attached and contractile podocytes.

These ultrastructural findings confirm that mechanical disequilibrium, specifically the concurrent loss of both shear stress and contractility, drives rapid podocyte detachment and barrier disruption, validating our model’s prediction that both orthogonal force components are required for maintaining the structural integrity of the filtration barrier.

### Stress-responsive integrin redistribution maintains adhesion under force imbalance

One of the key predictions from our simulation focused on the stability of integrins at different locations across the adhesion area of the foot process. In our model, the connection of actin to the slit diaphragm plays a critical role in maintaining the interconnection between nearby foot processes, while they are also connected to the integrin for stable anchoring. Consequently, different forces were calculated to act on integrins depending on their specific location when the system reached a state of equilibrium.

In low-stress situations, characterized by low shear stress and reduced actin contractility, the stability of integrin bonds at the very edge of the adhesion area appeared to be slightly higher than at other locations. As the stresses in the system increased, this difference between the edge and the center of the adhesion area became more pronounced (Fig. 3a, b). These findings predict accumulation of integrins in the most stable region, which corresponds to the periphery of the foot processes.

**Figure 3.**
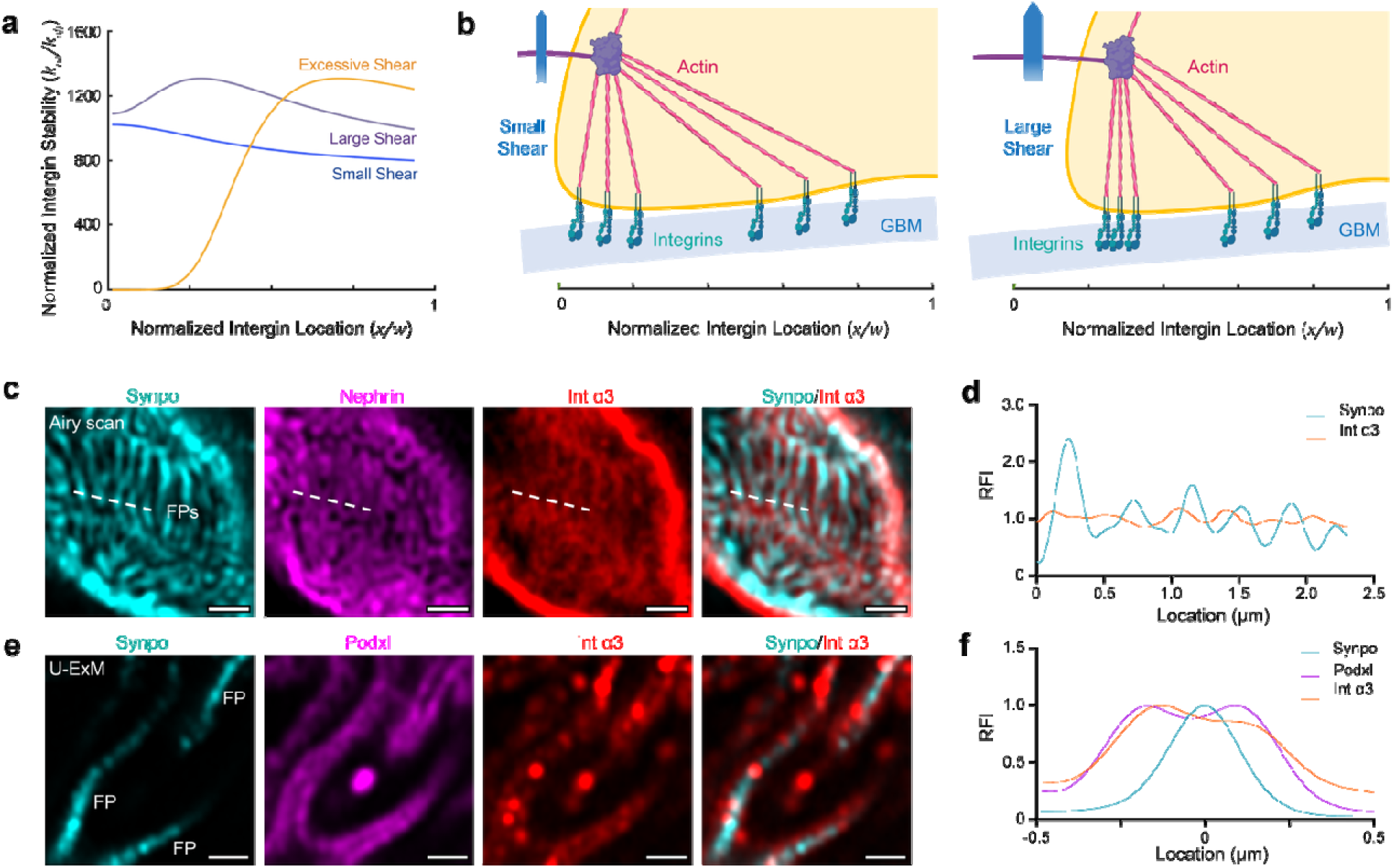
Mathematical modeling predicts stress-responsive integrin redistribution to foot process peripheries. **a,** Model-predicted integrin stability as a function of location from foot process periphery to center under varying shear stress conditions. Low stress (blue) shows minimal peripheral preference; increasing stress (orange to red) drives progressive peripheral accumulation with central depletion. **b,** Schematic of predicted integrin redistribution under mechanical stress. Integrins accumulate at foot process peripheries (green) as stress increases, with potential shape changes and edge detachment under excessive loading. **c,** Airyscan super-resolution imaging validates predicted pattern in healthy mouse glomerulus. Integrin α3 (red) accumulates in gaps between synaptopodin-marked foot processes (green), with nephrin marking slit diaphragms (blue). Scale bar: 1 μm. **d,** Relative fluorescence intensity (RFI) plot along indicated line in panel c shows integrin α3 peaks (red) localized between synaptopodin peaks (green), confirming peripheral accumulation pattern. **e**Expansion microscopy (4× expansion) enables single foot process resolution. Podocalyxin (membrane marker, magenta) encapsulates central synaptopodin (green), with integrin α3 (red) co-localizing at periphery. Scale bar: 1 μm. **f,** Quantitative analysis of straightened foot processes. Integrated RFI plot from all pixels surrounding foot processes shows central synaptopodin peak flanked by two peaks in both podocalyxin and integrin α3 channels, definitively confirming peripheral integrin localization matching model predictions.

To verify these predictions, we performed super-resolution imaging of the kidney glomerulus. Using Airyscan confocal imaging focused on the on-face area of the glomerulus, where foot process patterns can be identified through staining with synaptopodin (marking tense actin fibers) and nephrin (marking the slit diaphragm between foot processes), we observed an accumulation of integrin α3 signals, partially co-localizing with nephrin signals and surrounding the synaptopodin signals (Fig. 3c). Intensity maps along lines cutting through different foot processes showed a clearer pattern that integrin α3 accumulation was mostly observed between synaptopodin signals (Fig. 3d).

To achieve higher resolution, expansion microscopy was applied to get a clearer view of single foot processes. With this system, we could image foot processes at a resolution at which the center can be separated from the periphery though synaptopodin and podocalyxin marking. Integrin α3 localization around synaptopodin at the periphery of foot processes was clearly identified in most foot processes (Fig. 3e). Quantitative analysis, through straightening foot processes and measure fluorescence intensity changes based on their distance from the foot process center (Supplementary Fig. 4), revealed that straightened foot processes exhibited two peaks for both podocalyxin and integrin α3 surrounding the central synaptopodin peak (Fig. 3f), matching our prediction of integrin localization to the periphery.

When we examined integrin localization under different mechanical conditions, staining of integrin and synaptopodin revealed that integrin localized to the periphery of foot processes in both low and high blood pressure mice (Fig. 4a). However, intensity maps drawn along lines crossing multiple foot processes revealed that the intensity shifting in the high blood pressure mice was considerably larger than in low blood pressure mice (Fig. 4b), suggesting that accumulation of integrin was enhanced after the filtration rate increased.

**Figure 4.**
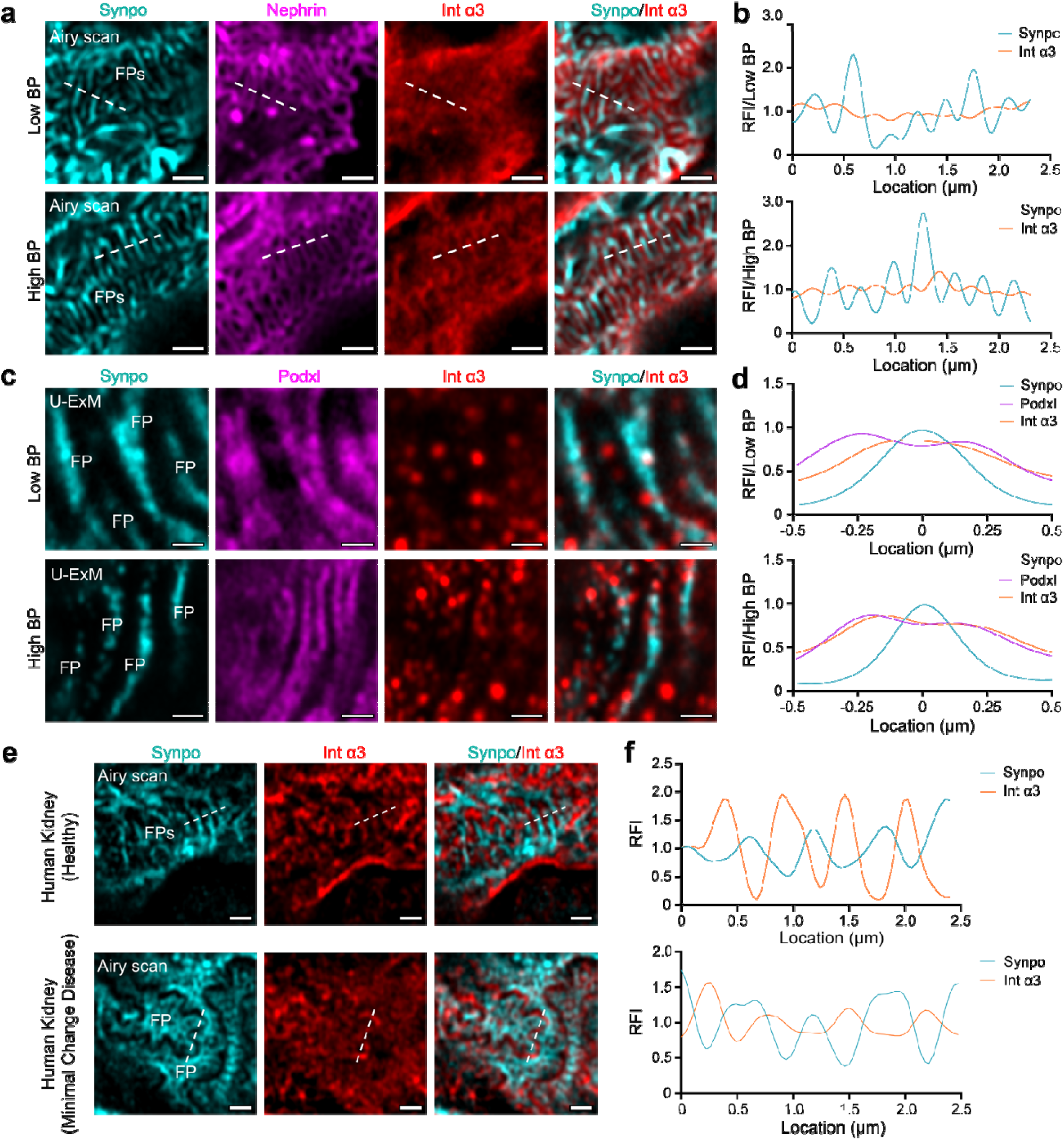
Mechanical stress alterations drive adaptive integrin redistribution in mouse models and in human kidney disease. **a,** Airyscan imaging reveals integrin α3 localization at foot process peripheries in both low BP and high BP mice 60 minutes post-blebbistatin. Synaptopodin marks central actin cables (green), integrin α3 shown in red. Scale bar: 1 μm. **b,** Relative fluorescence intensity plots along indicated lines in panel c demonstrate enhanced integrin accumulation in high BP mice. Greater peak-to-valley intensity differences in high BP samples indicate increased peripheral concentration under elevated shear stress. **c,** Expansion microscopy reveals foot process boundaries in both low and high BP mice. Podocalyxin staining (magenta) clearly identifies peripheries, with integrin α3 (red) accumulation partially lost in some low BP samples. Scale bar: 1 μm. **d,** Integrated RFI plots from straightened foot processes show differential integrin distribution. While podocalyxin maintains two peaks surrounding central synaptopodin in both groups, integrin α3 shows widened distribution in low GFR samples versus significant peripheral accumulation in high GFR group, confirming stress-dependent redistribution. **e,** Airyscan imaging of human kidney samples reveals conserved integrin localization patterns. In healthy human glomerulus (left), integrin α3 (red) localizes at foot process peripheries around synaptopodin-marked central actin (green). In minimal change disease (right), integrin α3 accumulates precisely between effaced foot processes, with sarcomere-like structures (SLSs) visible as discontinuous synaptopodin signals (arrows). Scale bar: 1 μm. **f,** Relative fluorescence intensity plots from human samples demonstrate peripheral integrin accumulation away from central synaptopodin signals in both healthy and diseased tissue, confirming conservation of the stress-responsive redistribution mechanism across species.

Further analysis using expansion microscopy showed that after averaging all imaged foot processes, synaptopodin followed accumulation in the foot process center, while podocalyxin exhibited two peaks at the periphery (Fig. 4c). For integrin intensity, although peaks at the foot process periphery were not clear after averaging, possibly due to non-symmetric distribution in most foot processes, a widened distribution could be identified along the width of the foot process in low GFR samples, while more significant accumulation to the periphery was noticed in the high GFR group (Fig. 4d), verifying our prediction that more accumulated integrin is needed under increased shear stress.

### Integrin redistribution patterns are conserved in human kidney disease

To establish the clinical relevance of our findings, we examined human kidney samples including both healthy kidney sections and sections from patients with minimal change disease, in which effaced foot processes are commonly found.^57^ These samples were subjected to similar staining procedures and Airyscan imaging to assess whether the integrin redistribution patterns observed in our mouse models are conserved in human kidney pathology.

Human kidney samples, including both healthy kidney sections and sections from patients with minimal change disease, revealed similar localization patterns of integrin α3 (Fig. 4e). Notably, in minimal change disease tissue where effaced foot processes were observed, integrin α3 signals were found perfectly localized between two nearby effaced foot processes, further validating our prediction regarding stress-responsive integrin redistribution as a mechanism to maintain adhesion under mechanical challenge.

The results revealed a similar localization pattern of integrin α3 compared to the findings in mouse tissue (Fig. 4e). Fluorescence intensity plots showed comparable patterns in both healthy and minimal change disease tissue samples (Fig. 4f), with integrin accumulation at the periphery of foot processes, away from central synaptopodin signals. This conservation of integrin localization patterns across species validates the fundamental nature of our proposed mechanical mechanism.

In the minimal change disease tissue, effaced foot processes could be observed while the integrin α3 signals were found perfectly localized between two nearby effaced foot processes (Fig. 4e). This clear accumulation of integrin α3 between effaced foot processes in minimal change disease tissues further validates our prediction regarding integrin localization and provides valuable insights into the response of podocytes to pathological conditions.

Moreover, within the effaced foot processes, we observed sarcomere-like structures (SLSs) that are known to be associated with increased stresses at focal adhesions during injury responses ^46^; these were evident from non-continuous, periodic synaptopodin signals (Fig. 4e). These structures are likely responsible for providing contractility, leading to increased stress on the integrins,^46^ and thus may contribute to the more obvious accumulation of integrin α3 around the periphery observed in this group. The presence of SLSs in diseased human tissue aligns with our model’s prediction that podocytes increase contractility to compensate for altered mechanical conditions during disease.

These findings highlight the significance of integrin localization and its potential implications for podocyte stability and function in health and disease. The conservation of stress-responsive integrin redistribution patterns from mouse models to human pathology suggests that the orthogonal force balance mechanism we identified represents a fundamental principle of podocyte mechanobiology with direct relevance to human kidney disease.

## Discussion

Our findings establish a fundamental mechanobiological principle governing cellular adhesion under competing mechanical forces. By revealing that podocyte attachment requires a homeostatic balance between external fluid shear stress and internal cellular contractility, we provide a mechanistic explanation for a longstanding clinical paradox: why reducing blood pressure alone fails to prevent podocyte loss in chronic kidney disease. The discovery that concurrent reduction of both mechanical inputs causes catastrophic barrier failure while individual reductions remain tolerable challenges current therapeutic approaches and suggests that maintaining mechanical homeostasis, rather than simply minimizing stress, should guide treatment strategies.

A key finding is that lowered blood pressure combined with reduced podocyte contractility causes more severe proteinuria than either perturbation alone. Hypertension is well-established as harmful to kidney health,^58^ and our experiments confirmed that elevated blood pressure causes proteinuria. Our results reveal the collaborative effects of all stresses in the system create unexpected vulnerabilities. Lowered blood pressure becomes particularly problematic when cell contractility is simultaneously compromised. This mechanical interdependence, demonstrated here for the first time, has important implications for understanding why some patients experience worsening kidney function despite blood pressure reduction and why angiotensin II blockade, which preserves podocyte contractility, proves more effective than blood pressure reduction alone ^16–18^.

Cell contractility emerges as a critical compensatory mechanism in our system. Recent work identified that podocytes form sarcomere-like structures (SLSs) in injured kidneys or during spreading in culture, suggesting a highly contractile phenotype ^46^. Our findings contextualize this observation: SLS formation likely represents an adaptive response where podocytes increase contractility to compensate for decreased shear stress from reduced GFR commonly observed after injury. The loss of SLSs in extended culture, which increases susceptibility to detachment ^46^, aligns with our model’s prediction that reduced contractility eliminates this compensatory capacity.

The stress-responsive redistribution of integrins to foot process peripheries represents a previously unrecognized adaptive mechanism. During injury, when foot process effacement occurs, the increased contractility evidenced by SLS formation drives exaggerated peripheral integrin accumulation. This redistribution pattern, conserved from mouse models to human minimal change disease, suggests an evolutionarily preserved mechanism for maintaining adhesion under mechanical challenge. The peripheral localization optimizes force transmission between the cytoskeleton and basement membrane, effectively creating a reinforced adhesive rim that resists detachment.

Our observation of rapid GBM thickening and undulation in certain area within one hour of contractility inhibition in low blood pressure mice reveals an unexpected role for podocyte mechanics in basement membrane organization. This finding connects to previous hypotheses that proteinuria relates to capillary dilatation from reduced compression forces ^59^. Our results provide direct evidence linking podocyte contractility to GBM morphology, suggesting that contractile forces along foot processes provide essential compression for maintaining basement membrane architecture.

The orthogonal force balance framework reconciles several puzzling clinical observations. Foot process effacement, common in most kidney diseases, reduces filtration area and paradoxically increases local shear stress when GFR is maintained. This explains why blood pressure reduction often helps: it temporarily reduces excessive shear on remaining foot processes. However, our findings suggest this approach may be counterproductive in diseases where podocyte contractility is already compromised, potentially explaining variable patient responses to antihypertensive therapy.

Our study has limitations. The acute blebbistatin model may not fully recapitulate chronic contractility loss in disease. The mathematical model, while predictive, simplifies the complex three-dimensional architecture of the filtration barrier. Future work should examine how these mechanical principles apply to specific kidney diseases with distinct patterns of mechanical dysfunction.

These insights open new directions for therapeutic intervention. Rather than universally lowering blood pressure, treatment could be tailored based on the mechanical status of podocytes. Biomarkers of podocyte contractility, such transient responses to changes in blood pressure, could guide whether to reduce pressure, enhance contractility, or both. Modulation of podocyte contractility without affecting systemic blood pressure might prove particularly valuable for patients with low blood pressure or compromised cardiac function.

## Conclusions

Our results establish an integrated mechanical force homeostasis as a fundamental requirement for maintaining biological barriers under stress. The identification of orthogonal force integration, where fluid and cytoskeletal mechanical inputs must be balanced to maintain cellular adhesion, represents a new principle in mechanobiology with implications beyond the kidney. Our findings challenge the prevailing paradigm that reducing mechanical stress universally benefits cellular health, demonstrating instead that cells require optimal mechanical loading from multiple directions to maintain structural integrity.

The conservation of stress-responsive integrin redistribution from mouse models to human disease validates this mechanism’s clinical relevance. By revealing how podocytes integrate competing mechanical forces to maintain the filtration barrier, we provide a mechanistic framework for understanding kidney disease progression and identify mechanical equilibrium as a therapeutic target. These principles likely extend to other barrier tissues facing competing mechanical forces, from pulmonary alveoli to the blood-brain barrier, suggesting that orthogonal force balance may be a broad requirement for maintaining tissue barriers under mechanical stress.

## Acknowledgments

Airyscan and TEM imaging was performed in the Washington University Center for Cellular Imaging (WUCCI). We thank Drs. Tyler Miller, MD and Eleanore Lederer, MD for reviewing the manuscript and for their valuable suggestions.

## Materials and Methods

### Mathematical Model of Podocyte Foot Process Adhesion

We developed a three-dimensional biomechanical model of podocyte foot process adhesion by adapting the molecular clutch framework^50,52^ to account for the unique orthogonal force environment of the glomerular filtration barrier. Unlike traditional models where forces act parallel to the adhesion plane, our model incorporated two mechanical challenges: vertical shear stress (*z*-direction) from glomerular filtration opposing horizontal contractile forces transmitted through actin cables connecting the cell body to foot processes.

#### Force Balance

The model considers force equilibrium at the junction where slit diaphragm connections meet central actin filaments. In the vertical direction:

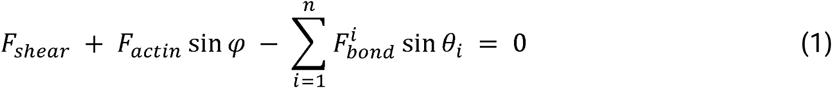

where *F_shear_* is the force arising from fluid shear stress, *F_actin_* is the upward force in the actin cable arising from actomyosin contractility in the cell body, φ represents the geometry of the foot process, *F_bond_^i^* is the force in one of the *n* cables connecting a slit diaphragm to focal adhesions, and *θ_i_* is the angle of that cable stretching towards the focal adhesion. At the glomerular basement membrane the total force *F_ECM_* resisted by the membrane is:

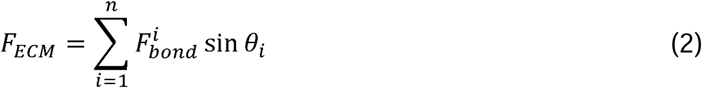

#### Kinetics

Individual integrin bonds experience force-dependent kinetics following:

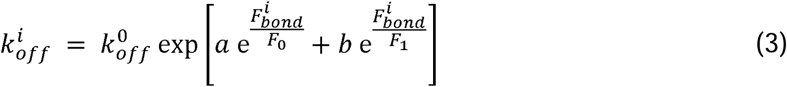

where parameters (a = -1.99, b = 0.723, *F_0_* = 0.0381 pN, *F_l_* = 0.0605 pN) were derived from force-dependent integrin αVβ3 bond lifetime data.^52^ The force on each bond is:

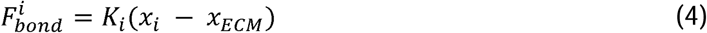

where *x_i_* is the displacement of connection *i, x_ECM_* is the displacement of the GBM, and *K_i_* is an effective spring constant that accounts for geometry:

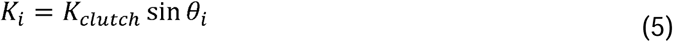

Myosin-driven actin retrograde flow velocity responded to mechanical loading according to:

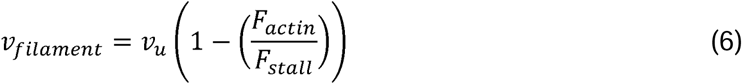

where *v_u_* is the unloaded velocity (baseline value 100 nm/s), *F_stall_* = *n_m_F_m_* is the total stall force from n_m_ myosin motors (baseline values *n_m_* = 800, *F_m_* = 2 pN).

#### Numerical Methods

Monte Carlo simulations were run using a stochastic framework with 10,000 time steps of duration Δ*t*=5 ms to reach steady state. A custom code was written using Matlab (The Mathworks, Natick, MA).

The algorithm began by assessing bond formation in each timestep, with unbound integrins forming bonds with probability,*p*_on_ =1 -exp(-k_on_Δ*t*), where the baseline value of k_on_ was 200 s^−1^.^52^ The displacement of the ECM was then assessed as

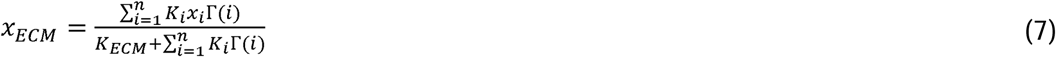

Forces on each bond were then determined from relative displacement, and bond breaking was assessed, with bonds breaking stochastically with probability *p_off_^i^* = 1 – exp(*k_off_^i^*Δt). Finally, positions were updated according to x_i_ (*t* + 1) = x_i_ (*t*) + *v_filament_Δt*.

Parameter space analyses were performed in which we systematically varied key parameters to map stability regimes. Shear stress was varied from 0-1000 pN; ECM stiffness (*K_ECM_*) from 0.001-100 pN/nm; cell contractility, via myosin motor number, from 100-1600 motors. Each simulation began with an initial integrin distribution of 75 bonds uniformly distributed across adhesion area.

System stability was assessed by the location of the maximum ratio of *k_on_/k_off_*, which was an indicator of integrin accumulation sites. Bond survival probability at steady state was also assessed, as was mean ECM displacement. Time to catastrophic detachment at which all bonds broke was also assessed.

#### Statistical Analysis

All simulations were run with multiple random seeds (*n* ≥ 10) to ensure reproducibility. Heatmaps were generated using Matlab with colormap scaling to visualize parameter dependencies. Convergence was verified by comparing steady-state values across different simulation durations.

### Blood pressure regulation in mice

Controlled increase of blood pressure was achieved through AAV-Renin vector injection, inducing Renin expression. The AAV-Renin vector was obtained from Charles Rivers (Houston, TX, USA). For induction of high blood pressure, 3 × 10^11^ of viral genomes were administered to each mouse through retro-orbital injection under anesthesia with isoflurane. To reduce blood pressure, a combination of anti-hypertension drugs, Losartan potassium (TCI, L0232, Tokyo, Japan) and Prazosin hydrochloride (Sigma, P7791, St. Louis, MO, USA), was administered to mice via the drinking water. 60 mg/100 mL Losartan and 10 mg/100 mL Prazosin were dissolved in the drinking water, which was refreshed every 2 weeks.

Blood pressure measurements were conducted as previously described ^60^. Briefly, a Millar pressure transducer (model SPR-671; Houston, TX) was inserted into the right common carotid artery of the mouse under continuous 1.5% isoflurane anesthesia. The blood pressure-time curve was recorded, and systolic and diastolic pressures were determined using the PowerLab data acquisition system (ADInstruments, Colorado Springs, CO, USA).

### Transdermal measurements for mouse GFR

GFR was measured transdermally using a monitor from Medi Beacon (St. Louis, MO, USA) and injection of FITC-sinistrin as previously described ^44^. Briefly, a small transdermal monitor was securely attached to the shaved back skin of each mouse, followed by the administration of FITC-sinistrin via retro-orbital injection. The fluorescent signal recorded by the monitor showed a rapid increase immediately after injection, followed by a continuous decrease as the FITC-sinistrin was cleared through glomerular filtration. The GFR of each mouse was calculated from the fluorescence intensity curve, with the rate of decrease representing the sinistrin clearance speed.

### Myosin inhibition in mice

The injection of the myosin inhibitor blebbistatin was performed as described ^61^. Briefly, 50 mM racemic Blebbistatin (Sigma, 203389, St. Louis, MO, USA) was prepared in a solvent of 25% Hydroxypropyl β-Cyclodextrin in PBS and injected intraperitoneally at 10 mg/kg using an injection volume of 10 ml/kg.

### Urine collection and creatinine and albumin detection

To assess the albuminuria levels, urine samples were collected before and after blebbistatin injection. Albumin, as a marker of kidney injury, was quantified and normalized to creatinine concentration to account for urine concentration variability. Creatinine concentration was measured using the QuantiChrom Creatinine Assay Kit (BioAssay Systems, Hayward, CA, USA) and used to determine the amount of urine to use for quantifying albumin. Albumin concentration was determined by running urine samples containing equivalent amounts of creatinine on a SDS PAGE gel followed by Coomassie blue staining. Standards with albumin concentrations ranging from 0.1 µg to 1.0 µg were run simultaneously to quantify the albumin in the urine samples using standard curve derived from the standards.

### Mouse kidney collection and fixation

For tissue collection, the mice were perfused with 4% PFA to remove blood cells and for initial fixation. Then, the whole kidney was excised, the capsule was carefully teased off, and the kidney was cut into two pieces and transferred to 4% PFA for fixation overnight. For paraffin embedding and sectioning, the kidney was dehydrated through an ethanol gradient and then into xylene and finally into melted paraffin. Paraffin sections of 5 µm were cut and picked up on charged glass slides for further analyses. For TEM imaging, slices of cortex were cut into 1 mm cubes, fixed in 2% glutaraldehyde, embedded in resin, ultra-thin sectioned, mounted on a specimen grid, and stained.

### Human kidney samples preparation

Formalin-fixed, paraffin-embedded nephrectomy specimens from a patient with renal cell carcinoma (non-tumor regions) obtained from King Chulalongkorn Memorial Hospital, Thailand, and renal biopsy specimens from a patient diagnosed with nephrotic syndrome due to minimal change disease from Barnes-Jewish Hospital, United States, were used to represent healthy and injured podocytes in vivo, respectively. The Ethics Committee of the Faculty of Medicine, Chulalongkorn University, approved the transfer and use of the sample from Thailand (Med Chula IRB no.0149/65) in compliance with international guidelines for human research protection, the Declaration of Helsinki and International Conference on Harmonization in Good Clinical Practice. The samples from Barnes-Jewish Hospital were obtained through Kidney Translational Research Center (KTRC). All participants provided written informed consent. The study was performed under conditions approved by the Institutional Review Board of Washington University School of Medicine.

### Immuno-fluorescent staining and imaging

For Airyscan imaging, immunofluorescence staining of kidney sections was performed based on a standard protocol. Briefly, paraffin-embedded kidney sections were rehydrated by passing through an ethanol gradient. Rehydrated sections were heated at 95 °C in PBS for 10 min for antigen recovery and submerged in 2% BSA for blocking. After 30 min of blocking, primary antibodies were diluted in 2% BSA and added to the sections for staining at 4°C overnight. Secondary antibodies diluted in 2% BSA were added to the sections after three PBS washes. After incubating for 1 h and washing in PBS, the samples were mounted for imaging.

The first and second antibodies used included: Guinea-pig anti-mouse synaptopodin (ARP, 03-GP94-N, Waltham, MA, USA); Rabbit anti-mouse integrin-α3 (BiCell, 10003, St. Louis, MO, USA); Goat anti-mouse nephrin (R&D System, AF3159, Minneapolis, MN, USA); Goat anti-mouse podocalyxin (R&D System, AF1556, Minneapolis, MN, USA); Alexa fluor-488 Donkey anti-guinea-pig secondary (Jackson ImmunoResearch, 706-545-148, West Grove, PA, USA); Alexa fluor-594 Donkey anti-rabbit secondary (Jackson ImmunoResearch, 711-585-152, West Grove, PA, USA); Dylight-405 Donkey anti-rabbit secondary(Jackson ImmunoResearch, 711-475-152, West Grove, PA, USA); and Alexa fluor-647 Donkey anti-goat secondary (Jackson ImmunoResearch, 705-605-003, West Grove, PA, USA).

Imaging was performed using a Zeiss 880 Airyscan platform. The system features a high-resolution galvo scanner along with two PMTs and a 32-element spectral detector as well as a transmitted light PMT for DIC imaging. In addition, the Airyscan unit provides sub-diffraction limited imaging down to 120 nm resolution. To visualize FP structure *en face*, a 60X objective with over 8 times zooming in was usually applied; FPs stained for synaptopodin could be identified from the SDs stained for nephrin.

### Sample embedding and expansion for super-resolution imaging

Imaging using expansion microscopy was conducted as described previously ^62^. Briefly, kidney tissues were collected, embedded as described, and sectioned for expansion microscopy. First, a mixture of PFA and acrylamide was added to the section to anchor the proteins to acrylamide. Next, acrylamide hydrogel crosslinking was performed on top of sections by an adding pre-crosslinking solution containing acrylamide, sodium acrylate, bis-acrylamide and the accelerator tetramethyl ethylenediamine (TEMED) and ammonium persulfate (APS).

The slides with hydrogel-embedded tissue samples were washed and submerged in a denaturing buffer containing SDS, NaCl, and Tris-Base, then heated in a 95°C water bath. Afterward, the hydrogels were collected from the slides and placed in fresh water for expansion, achieving approximately 4x expansion. The samples were then ready for immunostaining. After washing in PBS and blocking with BSA, the samples were treated consecutively with primary and secondary antibodies. Importantly, although PBS induced shrinking of the hydrogels during the staining process, the hydrogels were expanded again overnight before imaging. To further improve the resolution, the imaging of expanded kidney tissues was performed in the same Zeiss 880 Airyscan platform as detailed above.

### TEM imaging

Ultrathin sections from glutaraldehyde-fixed kidney pieces were imaged on a JEOL JEM-1400 120kV TEM system, which features a 0.38nm point resolution, 5-axis high-precision Piezo motorized stage suitable for large-area montages and AMT XR111 high-speed 4k x 2k pixel phosphor-scintillated 12-bit CCD camera, which features vibration-free Peltier cooling and 12µm pixel sizing. To visualize FPs, up to 4000X magnification was used.

### Membrane extraction and FIB-SIM imaging

FIB-SIM imaging on membrane extracted kidney samples were conducted as described previously ^51,63^. Briefly, isolated glomeruli were preincubated with extraction buffer containing 10 µM phalloidin, 10 µM Taxol, 2 mM ATP, 1 mM phenylmethylsulfonyl fluoride, 10 µM leupeptin in 0.1 M KCl, 30 mM HEPES at pH 7.2, 10 mM MgCI2, and 2 mM EGTA at pH 7.1, then subjected to extraction solution containing 0.01% saponin and 0.5% Triton X-100, and finally fixed with 4% PFA diluted in the extraction buffer before further fixed with 2% PFA/2% glutaraldehyde in PBS.

Samples were them prepared for FIB-SIM imaging through secondary fixation in 2% osmium tetroxide/1.5% potassium ferrocyanide in cacodylate buffer; staining in an aqueous solution of 1% thiocarbohydrazide; staining in aqueous 2% osmium tetroxide; staining in 1% uranyl acetate; *en bloc* stained for with 20 mM lead aspartate; dehydrating in a graded acetone series; and infiltrating into Durcupan resin. FIB-SEM imaging was conducted as described previously at a resolution of 10 nm/pixel with a dwell of 4 µs ^51^.

### Statistical Analysis

Significance analysis for foot process width was performed using one-way analysis of variance (ANOVA) followed by Tukey’s multiple comparisons test with the GraphPad Prism version 9.0 (GraphPad Software, San Diego, CA). Significance analysis for proteinuria evaluation was performed using Student’s t-test with GraphPad Prism version 9.0. For all data, differences were represented by significance based on the *P* value that was available upon each of the analyzed data. Data were expressed with each data point when available and together with means ± SD.

