## Supplementary Figures for "Orthogonal Force Balance Between Contractility and Shear Stress Governs Podocyte Dynamics"

Supplementary Figure 1

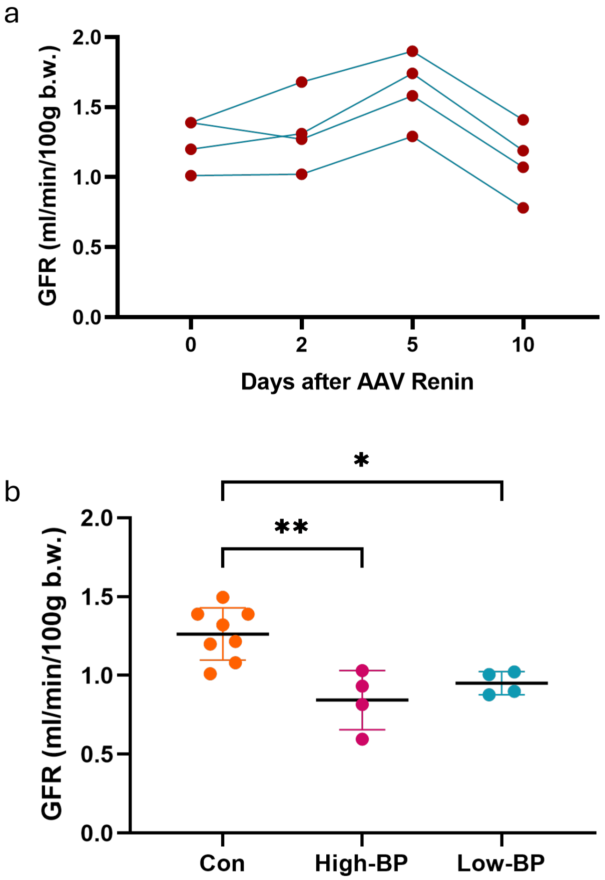


**Supplementary Figure 1, a,** AAV-Renin–induced hypertension leads to a gradual increase in glomerular filtration rate (GFR) within the first 5 days, whereas prolonged hypertension causes kidney injury and a reduction in GFR by day 10. **b,** One month after injection, GFR was reassessed, and both antihypertensive drug–treated mice and AAV-Renin–injected mice exhibited reduced GFR compared with controls.

Supplementary Figure 2

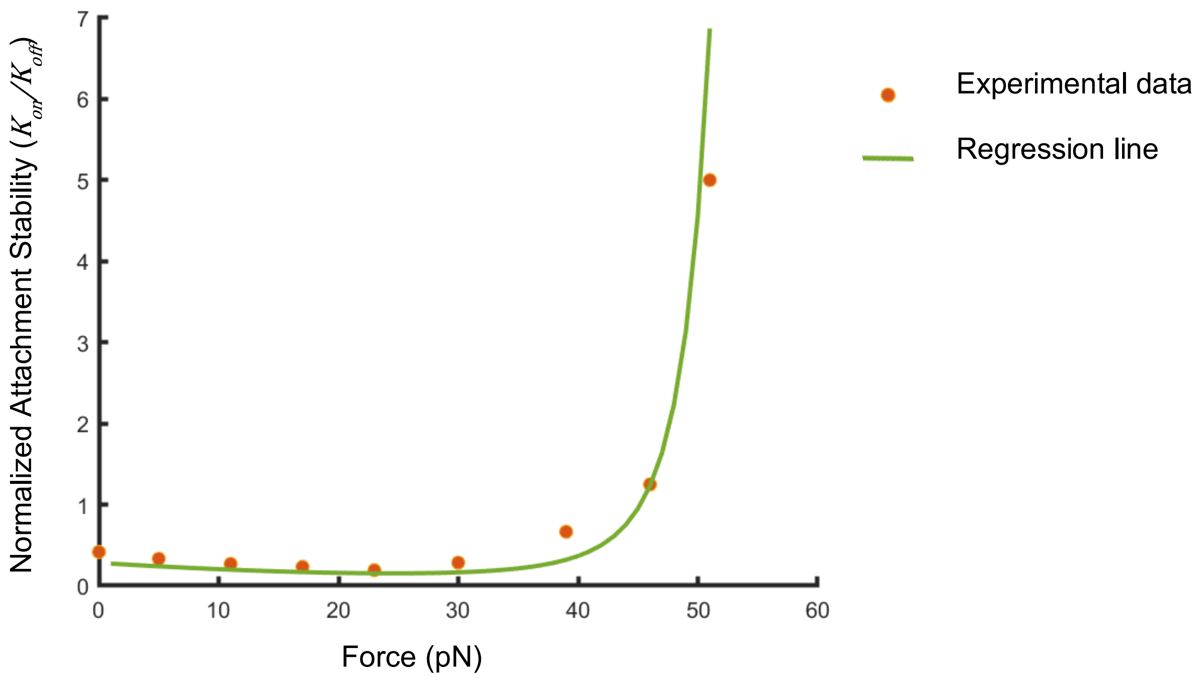


**Supplemental Figure 2,** Relationship between integrin αVβ3 stability and the force applied to the integrin. Experimental data were obtained from a previous study (Elosegui-Artola, A. et al. Nature Cell Biology **18**, 540-548 (2016)), and regression analysis was performed to fit the experimental data and predict integrin stability under varying force conditions.

Supplementary Figure 3

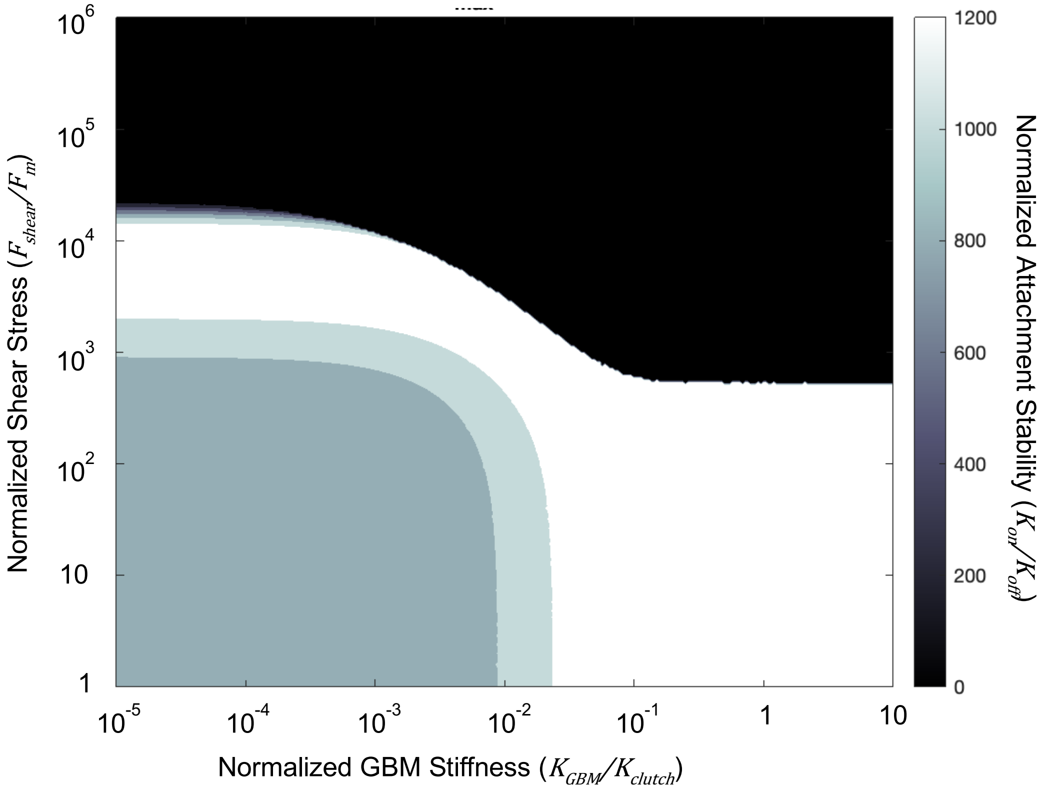


**Supplementary Figure 3,** Heat map illustrating how attachment stability is affected by varying fluid shear stress and glomerular basement membrane (GBM) stiffness. Attachment stability was predominantly influenced by shear stress. Increased GBM stiffness contributed positively to maintaining stable attachment under low shear stress, but negatively affected attachment stability under high shear stress.

Supplementary Figure 4

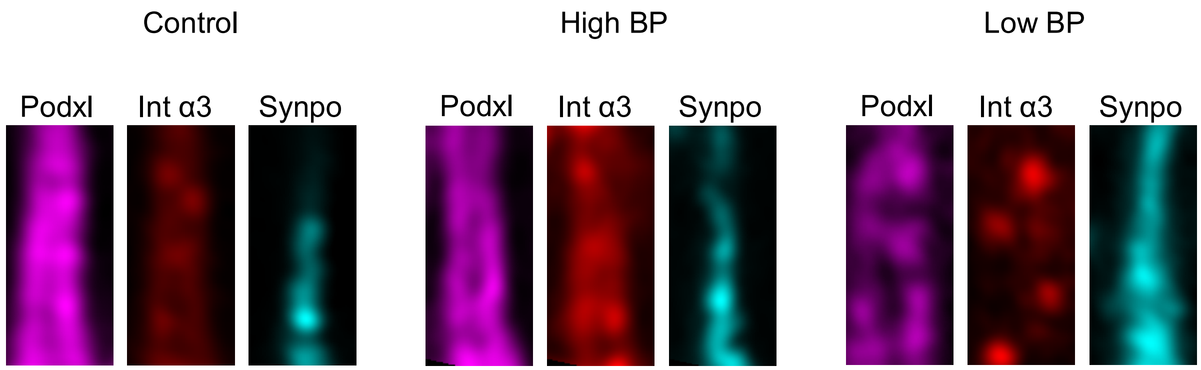


**Supplementary Figure 4,** Representative post-straightening foot process images acquired by expansion microscopy. Quantification was performed by calculating the mean fluorescence intensity as a function of distance from the central axis of the straightened image.
